# Social complexity during early development has long-term effects on neuroplasticity in the social decision-making network

**DOI:** 10.1101/2024.08.30.610458

**Authors:** Océane La Loggia, Diogo F. Antunes, Nadia Aubin-Horth, Barbara Taborsky

## Abstract

In social species, early social experience shapes the development of appropriate social behaviours during conspecific interactions referred to as social competence. However, the underlying neuronal mechanisms responsible for the acquisition of social competence are largely unknown. One key candidate to influence social competence is neuroplasticity, which functions to restructure neural networks in response to novel experiences or alterations of the environment. One important mediator of this restructuring is the neurotrophin BDNF, which is well conserved among vertebrates. We studied the highly social fish *Neolamprologus pulcher*, in which the impact of early social experience on social competence has been previously shown. We investigated experimentally how variation of the early social environment impacts markers of neuroplasticity by analysing the relative expression of the *bdnf* gene and its receptors *p75NTR* and *TrkB* across nodes of the Social Decision-Making Network. In fish raised in larger groups, *bdnf* and *TrkB* were upregulated in the anterior tuberal nucleus, compared to fish raised in smaller groups, while *TrkB* was downregulated and *bdnf* was upregulated in the lateral part of the dorsal telencephalon. In the preoptic area (POA), all three genes were upregulated in fish raised in large groups, suggesting that early social experiences might lead to changes of the neuronal connectivity in the POA. Our results highlight the importance of the early social experience in programming the constitutive expression of neuroplasticity markers, suggesting that the effects of early social experience on social competence might be due to changes in neuroplasticity.

## Introduction

During early ontogeny, the experienced social environment can lead to pervasive changes of phenotypic traits, including social behaviour (reviewed in Kasumovic & Brooks, 2011; Taborsky, 2017), dispersal decisions (Fischer et al., 2017; Gustafsson & Sutherland, 1988) and reproductive investment (Antunes & Taborsky, 2020; Naguib et al., 2006). Individuals which are socially deprived during development typically fail to develop the appropriate social skills later in life as shown in social spiders (Liedtke & Schneider, 2017), fish (Arnold & Taborsky, 2010; Fischer et al., 2015; Taborsky et al., 2012), mice (Branchi et al., 2006; D’Andrea et al., 2007), and apes (Kempes et al., 2008). Highly social species, such as cooperative breeders, typically engage in a diversity of interactions and form complex groups where individuals adopt different social roles and ranks in a mixed group of kin and non-kin (Clutton-Brock, 2006; Groenewoud et al., 2016a; Koenig & Dickinson, 2016; Taborsky, 2016). Consequently, they should particularly benefit from early acquisition of social competence, defined as the ability to optimally adjust one’s social behaviour to the prevailing social information, hence reducing potential costs associated to extended philopatry (Taborsky, 2021; Taborsky & Oliveira, 2012).

During early life, the Social Decision-Making Network (SDMN) is developing and different neural mechanisms, such as the monoaminergic and stress axis, are being adjusted (Antunes et al., 2021). The SDMN is a network of interconnected brain nodes responsible for processing and integrating social information and regulating the expression of social behaviour. The SDMN is well preserved across vertebrates (O’Connell & Hofmann, 2011, 2012). Early life experience impacts brain circuitry mediating defensive behaviour in rodents by, for example, modulating dendritic complexity and length in several brain regions of the SDMN, such as the hippocampus or the medial prefrontal cortex (Chocyk et al., 2013; Murthy & Gould, 2020; Soztutar et al., 2016). Social competence is thought to be accomplished by either rewiring or by biochemically switching nodes (i.e. through diffuse action of neuromodulators like neuropeptides, monoamines or hormones) of the SDMN (Cardoso et al., 2015). Neuroplasticity, which involves structural and functional changes in the brain, particularly through the influence of neurotrophins (Huang & Reichardt, 2001), plays a crucial role in mediating the rewiring of the SDMN and shaping social behaviour (Branchi et al., 2004).

Neuroplasticity functions to restructure neural networks in response to novel experiences or alterations in behaviour or environment (Kleim & Jones, 2008). Neurotrophins are secreted proteins that will act on cell-surface receptors of target cells, promoting survival and maturation of neurons (Bhattacharyya & Svendsen, 2003; Huang & Reichardt, 2001). Their activity mediates neuroplasticity in the short and long term, thereby facilitating effects of environmental experience on brain structure and function (Branchi et al., 2004). Therefore, neurotrophins are of key interest in the study of early social environmental effects on social competence. An important neurotrophin involved in the development of social behaviour and social competence of communally breeding mice is the brain-derived neurotrophic factor (BDNF) (Branchi et al., 2006), which is well conserved among vertebrates (Lucini et al., 2018). Thus, the BDNF pathways involving BDNF (brain-derived neurotrophic factor) itself and its two receptors are good candidates for investigating the long-term behavioural and neurological effects of early social experience in highly social vertebrates.

BDNF binds to the receptors p75NTR and TrkB (Huang & Reichardt, 2001; Purves et al., 2004), which have different functions in neuroplasticity. BDNF is known to be a major regulator of plasticity at excitatory synapses (Leal et al., 2015). There are two pathways for synaptic plasticity: BDNF/TrkB (Long Term Potentiation, long-lasting synaptic enhancement and synapse strength, i.e. the efficiency of the communication between two neurons at a synapse); and BDNF/p75NTR (Long Term Depression, synapse elimination) (Sakuragi et al., 2013). TrkB has a higher affinity for BDNF than p75NTR (Klein et al., 1991; Rodriguez-Tebar et al., 1990). The receptor TrkB has four functions, all related to synaptic plasticity: cell survival, neurite outgrowth, neuronal differentiation, and activity-dependent plasticity (Purves et al., 2004). TrkB plays a significant role in both social behaviour and stress vulnerability in mice (Razzoli et al., 2011). For instance, after repeated social defeat, mice expressing a truncated TrkB variant (leading to a decrease in BDNF signalling) exhibit more consistent social avoidance behaviours than their wild-type counterparts (Razzoli et al., 2011). In mice, *TrkB* knockout mutants perform poorly in complex and stressful learning tasks, suggesting that the TrkB receptor also has an important role in cognition (Minichiello et al., 1999). The receptor p75NTR has three main neuroplasticity functions: cell cycle arrest, cell death and neurite growth (Purves et al., 2004). In mice, downregulation of *p75NTR* in the hippocampus results in improved cognitive function (Maejima et al., 2018) and in autistic humans, *p75NTR* is correlated with impaired social cognition in a theory of mind test (Segura et al., 2015).

Experiments manipulating the early social environment in rodents have shown a short and long-term impact on brain neurotrophin expression, with an enriched social environment triggering higher expression (Branchi et al., 2006; Cirulli et al., 2003; D. Liu et al., 2000). In the cooperatively breeding fish *Neolamprologus pulcher*, early social deprivation and a social challenge interactively influence the expression of the *bdnf* gene in the hypothalamus, where fish reared in social deprivation show a downregulation of *bdnf* when not being challenged and an upregulation of *bdnf* when socially challenged by a dominant individual (Nyman et al., 2017). However, it is yet unknown (i) how *natural* variation of the early social environment impacts markers of neuroplasticity such as neurotrophins and their receptors and (ii) how differences in social competence are accompanied by differences in neuroplasticity. This will improve our understanding of the mechanisms underlying the development and expression of social behaviour.

The East African cichlid *N. pulcher* has emerged as a prominent model species for investigating the influence of the early social environment on neuroplasticity and social behaviour (Antunes & Taborsky, 2020; Arnold & Taborsky, 2010; Fischer et al., 2017; Nyman et al., 2017; Taborsky et al., 2012; Taborsky, 2016). These cooperative breeders form linear size-based hierarchies within groups, with social structures consisting of a dominant breeding pair and up to 25 subordinate helpers of various sizes and sexes (Dey et al., 2013; Groenewoud et al., 2016b; Taborsky, 2016). As cooperative breeders, these fish engage in numerous daily social interactions among group members, indicating the benefits of acquiring social competence early on (Taborsky et al., 2012).

In *N. pulcher* early-life environment affects the expression of social behaviour in that fish raised in larger, more complex social groups show more social competence than fish raised in smaller, less complex groups. This means they can more flexibly adjust their social behaviour to social information (La Loggia et al., 2024; Taborsky & Oliveira, 2012). We hypothesize that the ability to flexibly adjust one’s social behaviour is linked to neuroplasticity and therefore more complex early social environments should favour upregulation of synaptic plasticity mechanisms in the SDMN. Previous fish studies have shown that the social phenotype is linked to different *bdnf* expression patterns in the lateral zone of the dorsal telencephalic area (DL) (Teles et al., 2016) and in the hypothalamus (Nyman et al., 2017). Based on these studies, we expect that early social complexity influences *bdnf* expression patterns in the anterior tuberal nucleus (aTn), putative mammalian homologue: ventromedial hypothalamus VMH; in the lateral zone of the dorsal telencephalon (DL), putative mammalian homologue: hippocampus; in the preoptic area (POA), putative mammalian homologue: preoptic area plus paraventricular nucleus of the hypothalamus; and in the posterior tuberculum (TPp),putative mammalian homologue: ventral tegmental area VTA. The aTn and TPp are all involved in the expression of social behaviour (O’Connell & Hofmann, 2011) across vertebrates, therefore we expect to see different gene expression patterns to be related to early social complexity, with higher complexity being linked to the upregulation of synaptic plasticity pathways.

We investigated the relationship between early social experience and neuroplasticity in *N. pulcher.* We used individuals reared in two different levels of social complexity for their first 2 months of life (La Loggia et al., 2024) and analysed the relative brain gene expression of *bdnf* and its receptors (*p75NTR* and *TrkB*) across nodes of the SDMN. We performed brain microdissection of four brain nuclei of the SDMN all involved in the modulation of social behaviour (O’Connell & Hofmann, 2011) and used Quantitative real-time PCR to assess gene expression levels.

## Methods

### Rearing and housing conditions

We reared the fish in two early social conditions: large groups comprising ten adults (two breeders and eight unrelated subordinates), and small groups consisting of three adults (two breeders and one unrelated subordinate) (see details in La Loggia et al., 2024). In order to control for density of individuals within tanks we housed large groups in 300L-tanks and small groups in 100L-tanks. Fish were reared in their respective groups for 60 days after free swimming, before being housed in sibling groups in 20L tanks (same social conditions) during a ‘neutral’ phase of 60 days. At the age of 4 months the fish were tagged with coloured Visible Implant Elastomer (VIE) tags (Northwest Marine Technology Inc.) in the dorsal region to track individuals (Jungwirth et al., 2019). Behavioural tests were conducted between the ages of four months and one year to evaluate the effects of the early social environment on social behaviour (submission and aggression) and life history traits (exploration, helping and dispersal) (for detailed methods of these tests see La Loggia et al., 2024). After all behavioural tests were performed, i.e. when the fish reached 1 year of age, they were housed in sibling pairs in 50L tanks, separated from each other by clear, perforated partitions facilitating visual and olfactory communication. In total, we sampled 22 fish, eleven from large group-reared fish and eleven from small group-reared fish. As not all the siblings survived until sampling due to natural mortality, among the sampled fish there was a subset of individuals (three from each early-life treatment) that had been housed in 200L tanks alongside conspecifics from the same early life treatment.

### Candidate region of the SDMN

We investigated four nodes of the SDMN: (i) The anterior tuberal nucleus (aTn) is involved in aggression, reproduction and parental care in mammals (Félix & Oliveira, 2021; Lee et al., 2014; Y. Liu et al., 2019; McClellan et al., 2006; O’Connell & Hofmann, 2011; Olivier, 1977). In female plainfin midshipman fish *Porichthys notatus*, signs of neural activity are greater in the aTn when exposed to noises from conspecifics than ambient noise or heterospecific noises, suggesting that the aTn plays a role in the social behaviour network in fish (Mohr et al., 2018). (ii) The lateral zone of the dorsal telencephalic area (DL) is known to be involved in spatial learning via the storage of repeated experiences (Félix & Oliveira, 2021; O’Connell & Hofmann, 2011). In mammals, BDNF has been found to influence learning and memory in the hippocampus through its effects on long term potentiation (LTP) and long term depression (LTD) (Egan et al., 2003; Kovalchuk et al., 2002; Park & Poo, 2013). In zebrafish, BDNF involvement in the DL has been suggested to improve the ability to recognise dominant conspecifics (Teles et al., 2016). (iii) The preoptic area (POA) is involved in regulating sexual behaviour, aggression, and parental care in teleosts (Félix & Oliveira, 2021; O’Connell & Hofmann, 2011). (vi) The posterior tuberculum (TPp) is involved in reward-related behaviour like motivation and in reproductive social behaviours such as parental care (Félix & Oliveira, 2021; O’Connell & Hofmann, 2011; Trutti et al., 2019).

### Candidate genes

We measured expression of the three candidate genes: brain-derived neurotrophic factor (*bdnf)*, tyrosine kinase receptor B (*TrkB)* and nerve growth factor receptor b in teleosts, referred to as *p75NTR* here. For measuring *bdnf* expression, we used primers from Nyman et al., (2017). For measuring expression of *TrkB*, primers developed based on the *N. brichardi* mRNA sequence were used (NCBI database ID number XM_006787332.1, forward: GCTGGAACCACGATCCTCTG; reverse: GGGTCAGGTACACATTCTTGG, amplicon size 99), and for *P75NTR*, primers developed based on the *N. brichardi* mRNA sequence (NCBI database ID number XM_006807519.1, forward: GAATCAGGCACAAACAGTCGTCAAC; reverse: CTAAACAGCAGCTTCTCCACTTTCTC, amplicon size 93) were used. The expression of 18s was quantified as a house-keeping gene (Antunes et al., 2021; Nyman et al., 2017); we used primers from (Taborsky et al., 2013). Primers for *TrkB* and *P75NTR* were designed using primer-Blast (NCBI). The newly designed primers were tested prior to their utilisation. After performing a PCR (polymerase chain reaction) the products were sequenced and we run a Blast (NCBI) to confirm the specificity of the primers to the targeted genes.

### Tissue sampling

Tissue sampling was done as in Antunes et al., (2021). We euthanised the fish with an overdose of MS222 (Sigma-Aldrich). The decapitated heads were subsequently embedded in Tissue-Tek (optimal cutting temperature compound, OCT; Sakura) and frozen on dry ice within five minutes after euthanasia. The samples were stored and transported on dry ice until processing within the same day. We then sectioned the fish heads in the coronal plane, using disposable R35 microtome blades (Feather) on a Leica CM3050 cryostat. Slices were mounted on glass microscope slides immediately after being cut. The mounted slices were subsequently placed on a cold plate under a WILD M3C stereoscope, using a 24G sample corer tool (Fine Science Tools). The aTn, DL, POA and TPp were dissected. The collected brain tissue was stored for each individual per brain region in an Eppendorf 1.5ml tube with 100µl of DNA/RNA Shield (Zymo Research), and then stored at -80°C until further processing.

### RNA extraction

Brain tissues were digested in Proteinase K for 2h at 55°C to lyse the tissue before RNA extraction. We extracted RNA following the protocol from the Quick-RNA MicroPrep kit (Zymo Research). Samples were treated with DNase I (Zymo Research) to avoid DNA contamination. Extracted RNA was quantified using QuBit RNA HS assay kit (ThermoFischer Scientific) on a QuBit 2.0 fluorometer machine (ThermoFischer Scientific; sample RNA concentration ranged from 10.6 to 70.2 ng µl^−1^). RNA concentration was too low to be detected in 32 cases out of 88 samples. We reverse transcribed all samples to cDNA using an iSCRIPT cDNA synthesis kit (Bio-Rad).

### Quantitative real-time PCR

Quantitative real-time PCR (q-rt-PCR) and melting curves (ranging from 50 to 90°C) were conducted in triplicate using standard curves for 5 × 10-fold dilutions of all brain RNA. For the *p75NTR* primers, we used 5 × 10-fold dilutions of gBlocks® gene fragment synthesised from *p75NTR* predicted sequence from *N. brichardi*. These analyses aimed to assess the amplification efficiency (E) of each primer pair, as well as to ensure the absence of primer dimers and the specificity of the amplification (Antunes et al., 2021; Aubin-Horth et al., 2012). The primers (Microsynth) and 1 µl of sample cDNA were prepared in a 96-well plate (Greiner Bio-one). We added in each plate 5× HOT FIREPol EvaGreen qPCR Mix Plus ROX (Solis BioDyne) and performed q-rt-PCR on an ABI PRISM 7000 (Applied Biosystems). Triplicate runs were performed for all cDNA samples, including no-template controls. Melting curves were conducted for each replicate to confirm the absence of primer dimers and ensure the production of a single-amplified product. Cycle thresholds (Ct) were determined for each sample, and gene expression for individual brains was calculated using the formula 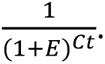 The relative expression was then normalized to the reference gene (18s) (Pfaffl, 2001).

### Statistical analysis

Statistical analyses were conducted using R software version 4.2.2 (R Core Team, 2020). In certain cases, gene expression data for specific genes were excluded from the analysis due to a high coefficient of variation (CV) among the three replicates. To ensure data quality, samples with a CV exceeding the predetermined cut-off of 5% were removed from the analysis. We analysed the effect of early social environment (small vs large groups) on expression of each gene within each node of the SDMN by fitting linear models (LMs). All LMs initially included sex, early-life experience, exposure to behavioural tests and age at sampling as fixed factors. Through stepwise model reduction, factors with an Akaike information criterion (AIC) difference greater than 2 were systematically removed from the models and are not presented in the table. However, early life experience was consistently retained in all models, as it was the experimental factor in this study addressing the research question. To enhance the robustness of the analysis, outliers were identified and removed by calculating Cook’s distance. This approach allowed the detection of influential data points, which were subsequently excluded from the analysis. By removing outliers, we ensured that the statistical models were not unduly influenced by extreme observations, resulting in more robust and accurate results. Twenty outliers were removed out of 264 data points: 8 from the aTn data points; 5 from the DL data points; 1 from the POA data points and 6 from the TPp data points. Normality assumptions for the error term were evaluated using Shapiro-Wilk tests and visual inspection of quantile-quantile plots for skewness and kurtosis. Homogeneity of variance was assessed through Tukey-Anscombe plots. To meet normality assumptions, gene expression levels were log-transformed or Box-Cox transformed.

## Results

The results revealed significant differences in gene expression between fish raised in large vs. small groups across different brain regions. In the anterior tuberal nucleus (aTn), fish raised in large groups exhibited an upregulation of *bdnf* and *TrkB* compared to fish raised in small groups (Table 1a, Figure 1). In the dorsal telencephalon (DL), fish raised in large groups displayed an upregulation of *bdnf* and a downregulation of *TrkB* compared to fish raised in small groups (Table 1b, Figure 2). Furthermore, in the preoptic area (POA), fish raised in large groups showed an upregulation of *bdnf*, TrkB, and *p75NTR* compared to fish raised in small groups (Table 1c, Figure 3). In the posterior tuberculum (TPp), we found no effect of early-life environment on gene expression for the three investigated genes (Table 1d, Figure 4). *bdnf* expression increased with age in the aTn, DL and POA, whereas it decreased with age in the TPp (Table 1a, Table 1b, Table 1c). *p75NTR* increased with age in the POA (Table 1b). *TrkB* increased with age in the aTn and decreased with age in the TPp (Table 1d). Sex influenced *TrkB* expression in the aTn (Table 1a), *bdnf* expression in the DL (Table 1b) and *TrkB* expression in the TPp (Table 1d).

**Figure 1:**
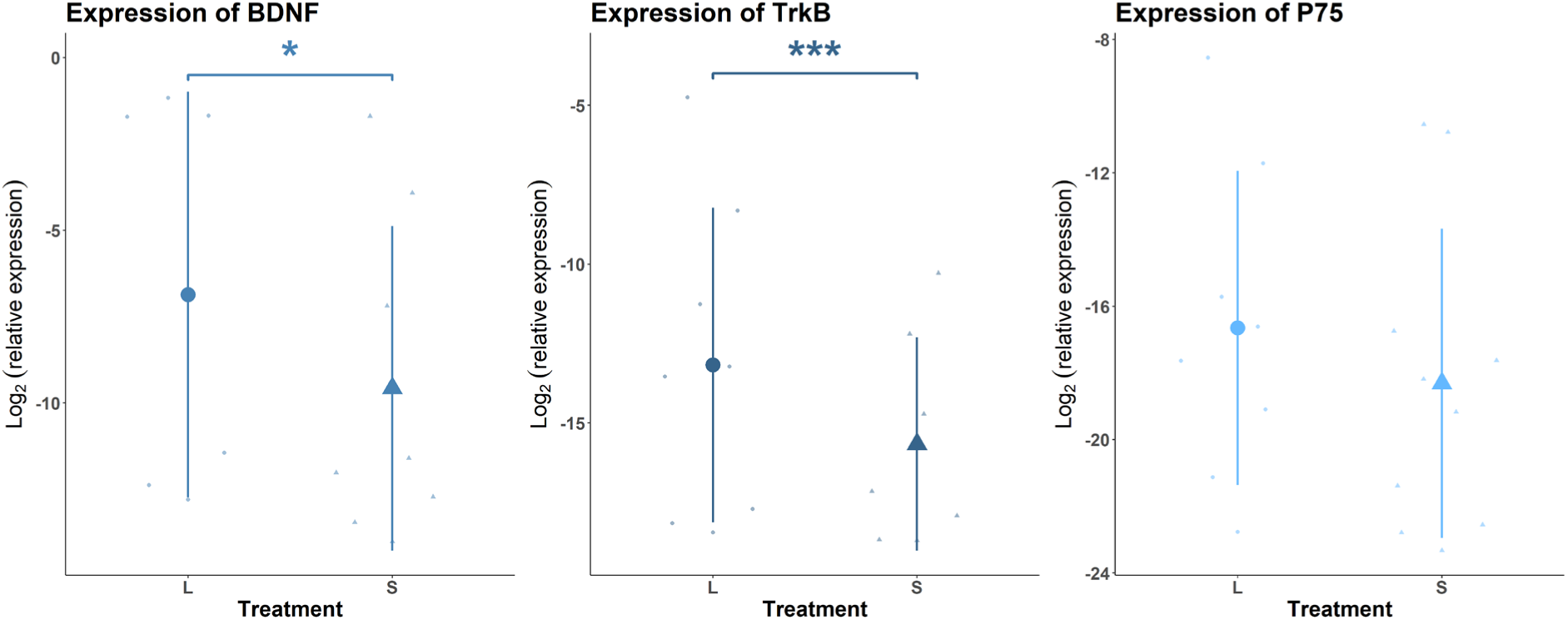
Relative expression of the three candidate genes in the ATN. L is for fish raised in large groups and S for fish raised in small groups. Bars represent standard error and dots means. Significance bars based on model output: * p<0.05, ** p<0.001, *** p<0.0001.

**Figure 2:**
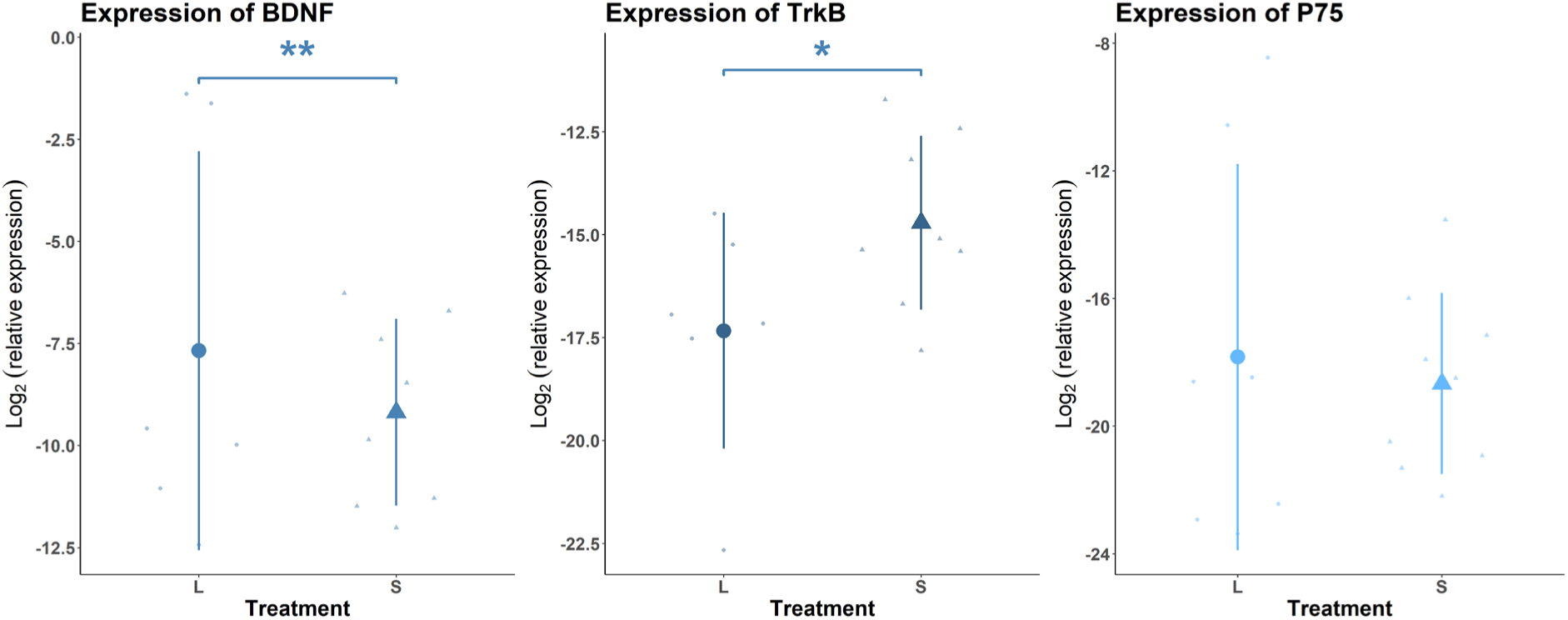
Plot of relative expression of the three candidate genes in the DL. L is for fish raised in large groups and S for fish raised in small groups. Bars represent standard error and dots means. Significance bars based on model output: * p<0.05, ** p<0.001, *** p<0.0001.

**Figure 3:**
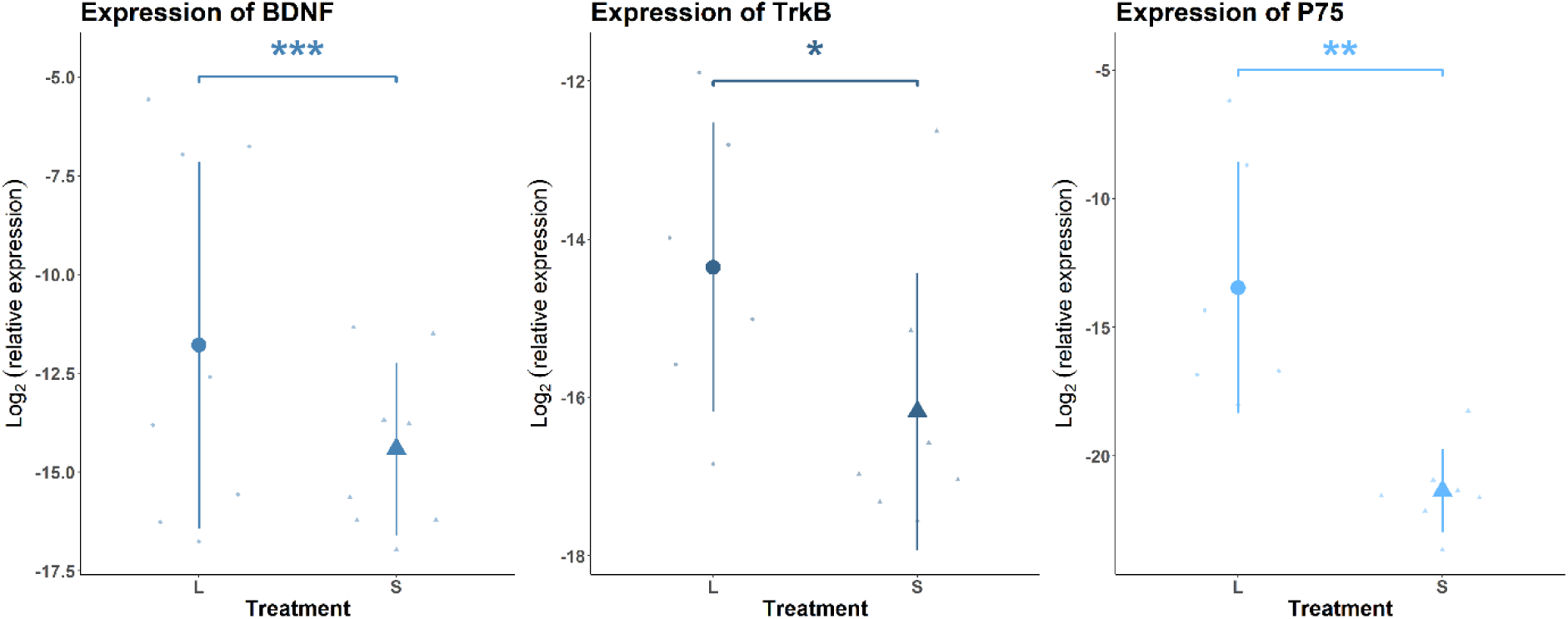
Plot of relative expression of the three candidate genes in the POA. L is for fish raised in large groups and S for fish raised in small groups. Bars represent standard error and dots means. Significance bars based on model output: * p<0.05, ** p<0.001, *** p<0.0001.

**Figure 4:**
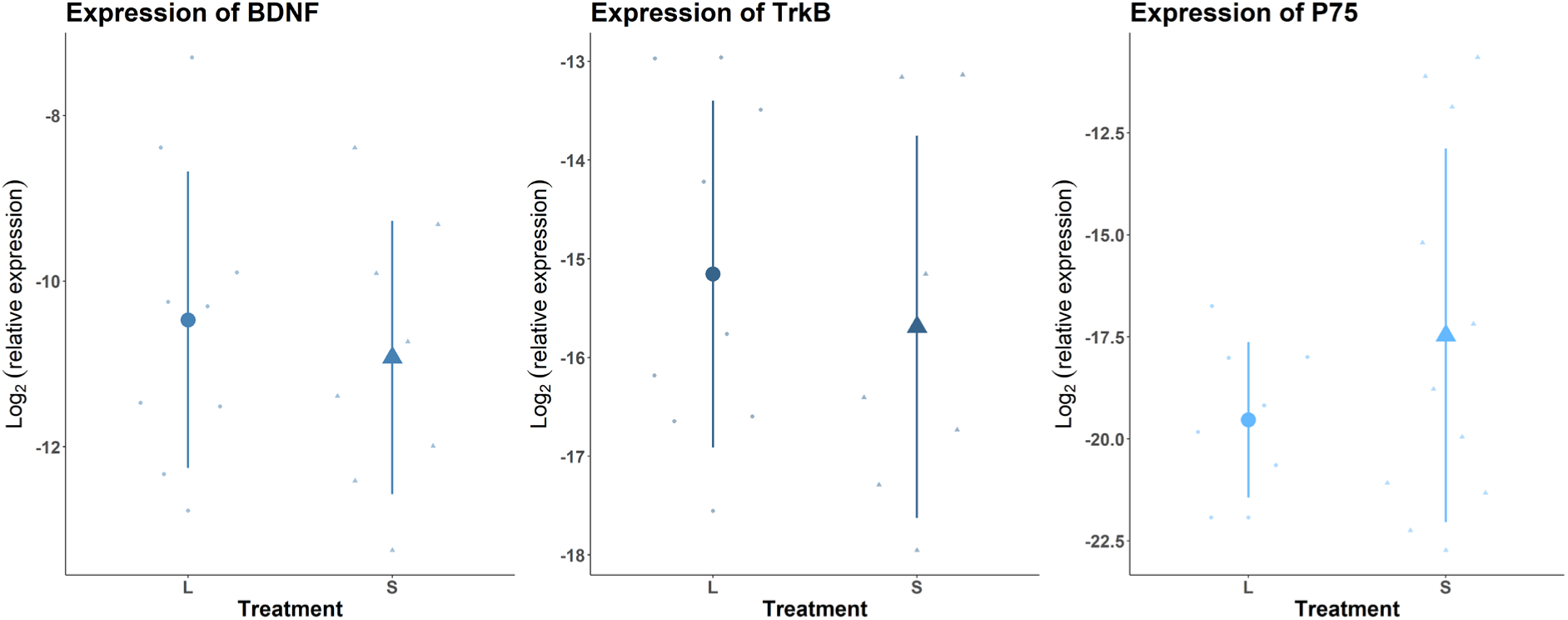
Plot of relative expression of the three candidate genes in the TPp. L is for fish raised in large groups and S for fish raised in small groups. Bars represent standard error and dots means. Significance bars based on model output: * p<0.05, ** p<0.001, *** p<0.0001.

**Table 1:**
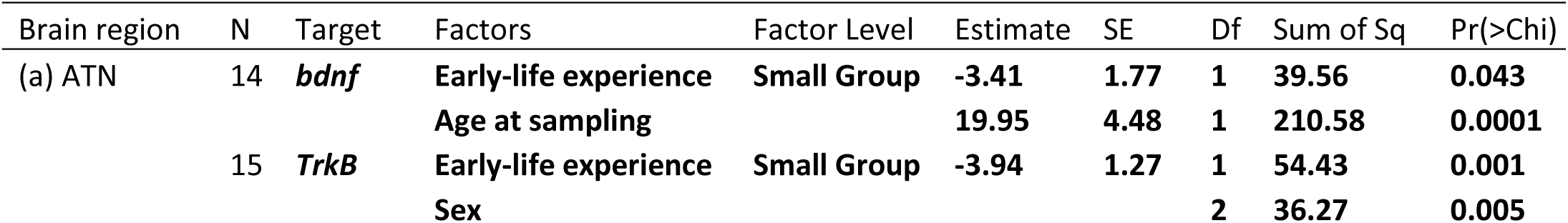
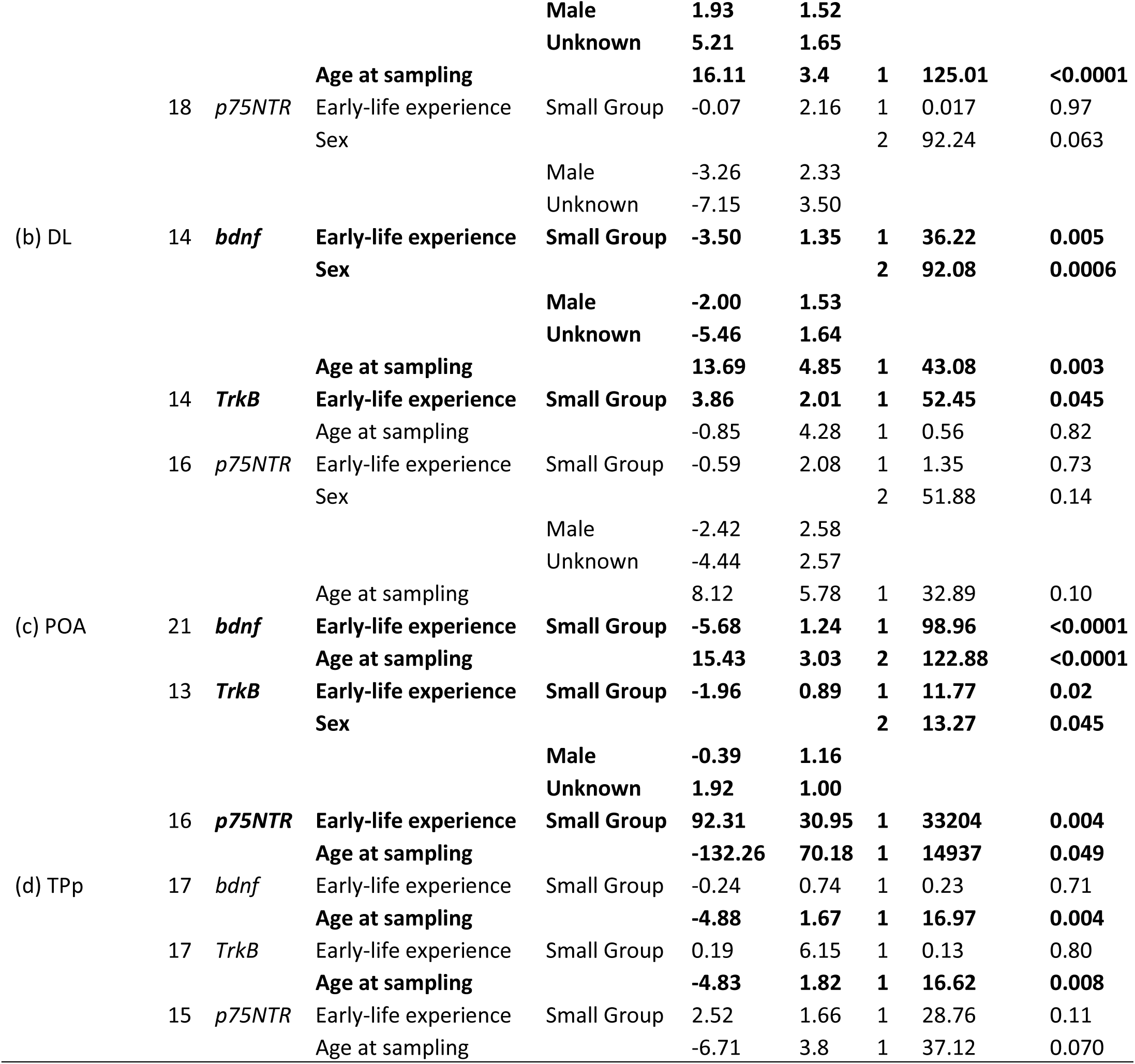
Results from linear models on the effects of early social experience on the relative expression (log2) of three candidate genes in (a) the ATN, (b) the DL, (c) the POA and (d) the TPp. In model (c) for the p75NTR gene, relative gene expression was Box–Cox transformed to achieve normality of model residuals; the interpretation of estimates has to take account of this data transformation, which caused the estimate signs to reverse in direction. Significant p-values are highlighted in bold.

## Discussion

We investigated the effect of the early social environment on neuroplasticity markers in four nodes of the SDMN. We demonstrated that the early social environment induced long-term differences in expression of *bdnf* and its receptors in three of the four nodes of the SDMN. There was an upregulation of both *bdnf* and *TrkB* in the aTn of fish raised in the larger groups, suggesting that fish raised in large groups might use the BDNF/TrkB pathway more in the aTn compared to those raised in small groups. In the DL, the putative homologue of the hippocampus, *TrkB* was downregulated in large groups, while *bdnf* was upregulated in these groups. In the POA, all three genes were upregulated in fish raised in large groups compared to fish raised in small groups. Our results demonstrate that social complexity during early development has long-term effects on neuroplasticity.

The BDNF/TrkB pathway promotes neuronal survival and differentiation, as well as neurite growth (Park & Poo, 2013; Purves et al., 2004). The BDNF/TrkB pathway has been denoted as the synaptic enhancement pathway (Sakuragi et al., 2013). In contrast, the BDNF/p75NTR pathway induces apoptosis, decreases neuron survival but also promote neurite growth (Park & Poo, 2013; Purves et al., 2004). The BDNF/p75NTR pathway has been denoted as the synaptic elimination pathway (Sakuragi et al., 2013).

The upregulation of the BDNF/*TrkB* pathway in the aTn of large-group fish could be an indication that enriched early social experiences might lead to an increased number of synapses and dendrites and the presence of more neurons in this brain area (Park & Poo, 2013; Purves et al., 2004; Sakuragi et al., 2013). In rodents, the VMH (putative homologue of the aTn) plays a critical role in regulating social fear and fear of predators (Silva et al., 2013). Fish raised in large group may benefit from more safety due to the presence of more individuals to guard the juveniles, as a result juveniles can engage in more social activities (Arnold & Taborsky, 2010; Fischer et al., 2015). Prior results in *N. pulcher* have demonstrated that predator fear response is mediated by early social experience. Fish raised in presence of parents display decreased fear response towards predators (Watve & Taborsky, 2019). We hypothesise that the upregulation of the BDNF/TrkB pathway in the aTn might be due to the decrease of intrinsic fear induced by exposure to an enriched early social environment, which might have led to the increase of synapses and neurons in the aTn. This hypothesis is partially supported by previous results showing that fish raised in larger group were found to have a larger hypothalamus (Fischer et al., 2015)

In the DL, *TrkB* was downregulated and *bdnf* was upregulated in fish raised in large groups. Blockage of BDNF/TrkB pathway was found to impair long term potentiation (LTP) in the hippocampus of mammals (Park & Poo, 2013). In-vitro experiments showed that masking of *TrkB* in the hippocampus of mice induce an alternative activation of p75NTR by BDNF (Sakuragi et al., 2013). This may suggest that in the DL, large group fish are more likely to use the synaptic elimination pathway and are therefore more capable of rewiring circuits in response to novel cues from the environment. Moreover, expression of *bdnf* in the DL of zebrafish was suggested to modulate social memory, as it improved the recognition and memory of dominant conspecifics (Teles et al., 2016). Taken together, this suggests that fish raised in larger groups can display higher sensitivity to environmental stimuli.

In the POA of fish raised in large groups, both receptors and *bdnf* were upregulated. This could mean that both pathways are acting in synergy. The BDNF/p75NTR pathway might ensure neuron survival, while the BDNF/TrkB pathway promotes increased connectivity between neurons (Sakuragi et al., 2013; Zanin et al., 2019). As shown in the mammalian hippocampus, TrkB and p75NTR interact after BDNF attaches to TrkB, and this interaction is important for activating a specific cell survival pathway (Zanin et al., 2019), suggesting that this could happen as well in our study in the POA. The development of the POA is known for its sensitivity to early life social experience. In guinea pigs, social instability during early-life causes an upregulation of oestrogen and androgen receptors in the POA (Kaiser et al., 2003). Additionally, early-social deprivation has life-long effects on the POA’s neurogenomic profile (Antunes et al., 2021). In the present study, we show that early social experience also leads to long-lasting effects in neuroplasticity in the POA, which might have led to an increase in neuronal connectivity in this region. Such constitutive changes in neuronal plasticity, possibly affecting long-term changes in neurogenomic profiles, could be responsible for the observed long lasting effects of social competence (La Loggia & Taborsky, 2024).

The four brain areas targeted in our study are linked within the SDMN (O’Connell & Hofmann, 2011). The POA is mutually interconnected with DL and aTn, whereas aTn and DL are interconnected, and so are DL and TPp (O’Connell & Hofmann, 2011). In human children, BDNF was found to be of particular importance in modulating the connectivity between brain regions. Carrier of a BDNF gene variant showed different patterns of connectivity within the hippocampus and between the amygdala, insula and striatal regions (Thomason et al., 2009). Due to the connectivity between our candidate regions, we speculate that the BDNF pathway plays an important role in facilitating information transfer not only within but also between these regions.

Early social experience is known to influence social competence in animals. Social competence is based on behavioural flexibility (Oliveira, 2009; Taborsky & Oliveira, 2012), which might be regulated via neuroplasticity. We showed previously that enriching the early social environment by enlarging group size enhances social competence (La Loggia et al 2024). Using the same batch of fish as La Loggia et al. (2024), here we report multiple effects of early social experience on the expression of *bdnf* and the genes of its receptors. Taking the results of the previous and current study together, it suggests that the effect of social experience on social competence might be mediated by BDNF and its receptors, as we had hypothesized. In line with our findings, communal nesting experience in early life of mice led to higher rates of displayed maternal care later in life, as well as higher expression of neurotrophins in the brain (Branchi et al., 2006). In *N. pulcher*, the early life family-group composition influenced the expression of the *bdnf* gene in the hypothalamus (Nyman et al., 2017).

Cardoso and colleagues (2015) suggested that at the neuronal level, social competence can be mediated by two mechanisms, neural structural reorganisation or a biochemical activation of neuronal circuits. It has been shown that neurotrophins are key in mediating neuro-behavioural plasticity (Branchi et al., 2004). We hypothesise that structural reorganisation might be mediated by the BDNF/p75NTR pathway, where cell death could lead to restructuring of the neural circuits and therefore help flexibly reallocating space or neural resources in order to cope with environmental changes. On the other hand, biochemical activation of relevant neural circuits might be achieved through the BDNF/TRKB pathway, with more neurons and/or more connections between neurons facilitating the change of neural circuits.

A likely mechanism regulating the observed long-term effects on social competence is learning, either social learning or learning by experience (Taborsky et al., 2016). Repeated social experience might lead to learning and help consolidating long-term memory (Gower, 1990; Isabel et al., 2004). Neurotrophin production is associated with learning. In rodents, contextual fear conditioning leads to an increase in *bdnf* expression in the hippocampus (Chen et al., 2007). In *N.pulcher*, during early ontogeny individuals are exposed to repeated interactions with multiple partners, which might lead to learning how to appropriately behave in social interactions. It is possible that different early-life experience leads to different neural activity during the experience phase, resulting in different constitutive expression of neurotrophins such as *bdnf* (Branchi et al., 2004; Park & Poo, 2013). Long-term potentiation (LTP) and long-term depression (LTD) are mechanisms underlying learning and the formation of memory (Sakuragi et al., 2013). Repeated LTP induction leads to long-lasting synaptic enhancement coupled with synaptogenesis (Tominaga-Yoshino et al., 2002, 2008). This is mediated by the BDNF/TrkB pathway (Sakuragi et al., 2013). On the contrary, repetitive LTD induction leads to long-lasting synaptic suppression coupled with synapse elimination (Kamikubo et al., 2006; Shinoda et al., 2005), which is mediated by the BDNF/p75NTR pathway (Sakuragi et al., 2013). We hypothesise that large groups face more opportunities to experience repeated social interactions with multiple different individuals by the presence of more adults interacting with each other in different ways. Therefore, fish raised in larger, more complex groups should be more likely to experience repeated LTP or LTD, mediated by BDNF/TrkB and BDNF/P75NTR respectively. This hypothesis should be verified experimentally, by investigating the formation of LTD and LTP during early-life within the brain regions investigated here.

While we showed early-life experiences influenced neurotrophin expression, it is important to note that neurotrophin expression could also be influenced by the current environment. A previous study showed that in *N. pulcher* the differential expression of *bdnf* in the hypothalamus resulted not only from early experience, but was interactively influenced also by a recent social challenge (Nyman et al., 2017). Moreover,, in zebrafish *bdnf* expression depended on the outcome of a contest (Teles et al., 2016). In this study, we controlled for the current environment by isolating all individuals 24h prior to sampling. Hence, our results cannot be explained by *bdnf* expression in response to current challenges. Therefore, our results indicate constitutive differences in neuroplasticity that can only be explained by the experimental differences in early-environment treatments.

Age and sex influenced relative gene expression in different regions. Sex influenced the expression of *bdnf* in the DL, the expression of *TrkB* in the aTn and the POA and the expression of p75NTR in the aTn and in the DL. Due to our lack of power (six fish were too young to determine their sex) we cannot interpret the direction of the differences in gene expression between sexes. Even though our fish differed in age by less than three months (age on average 3.44 ± 0.22 years), we observed an age effect on *bdnf* expression. Older fish showed an increased expression of *bdnf* in the aTn, DL and POA but a decreased expression in the TPp. In the DL older fish had higher expression of *p75NTR*. Finally older fish showed higher expression of *TrkB* in the aTn but lower in the TPp. In rodents, expression of *bdnf* and *TrkB* is lost in the prefrontal cortex of aged individuals (Coria-Lucero et al., 2016). Our results suggest that even small age differences lead to differences in the expression of *bdnf* and its receptors that seem to be region specific, and thus age and sex should be considered as factors in future gene expression experiments.

In conclusion, our results highlight the importance of early social experience in programming the constitutive expression of neuroplasticity markers. Here we showed that the expression of two distinct pathways regulating neuronal plasticity are shaped by early social experiences which persist into adulthood; and previously we showed that the same early social experience modulate social competence (La Loggia et al., 2024). Taken together, we hypothesize that early social experiences alter social competence via the BDNF/TrkB and the BDNF/p75NTR pathways. Future work should aim to establish the causal link between the neurotrophin pathways and social competence in animals by manipulating the activity of the two pathways. Generally, more research is needed to understand the mechanisms underlying the expression of social competence in social animal systems.

## Acknowledgements

We thank Maria Reyes for help with maintenance of the family groups. Magda Teles for valuable input on the *N. pulcher* brain atlas. Evi Zwygart, Danielle Bonfils and Markus Wyman for animal care and technical support; and the entire Hasli team for discussions.

## Authors’ contributions

O.L.L.: conceptualization, data curation, formal analysis, investigation, methodology, writing - original draft, writing-review and editing; D.F.A..: conceptualization, formal analysis, investigation, methodology, writing - original draft, writing - review and editing; N.A.H.: conceptualization, supervision, writing - review and editing, B.T.: conceptualization, funding acquisition, project administration, supervision, writing - review and editing.

## Data availability

Data is available on DataDryad. (For review as supplement)

## Benefit-Sharing Statement

Benefits from this research comes from sharing our data and results in public databases (see data availability statement).

## References

Antunes, D. F., & Taborsky, B. (2020). Early social and ecological experience triggers divergent reproductive investment strategies in a cooperative breeder. Scientific Reports, 10(1), 1–8. 10.1038/s41598-020-67294-x

Antunes, D. F., Teles, M. C., Zuelling, M., Friesen, C. N., Oliveira, R. F., Aubin-Horth, N., & Taborsky, B. (2021). Early social deprivation shapes neuronal programming of the social decision-making network in a cooperatively breeding fish. Molecular Ecology, 30(16), 4118–4132. 10.1111/mec.16019

Arnold, C., & Taborsky, B. (2010). Social experience in early ontogeny has lasting effects on social skills in cooperatively breeding cichlids. Animal Behaviour, 79(3), 621–630. 10.1016/j.anbehav.2009.12.008

Aubin-Horth, N., Deschênes, M., & Cloutier, S. (2012). Natural variation in the molecular stress network correlates with a behavioural syndrome. Hormones and Behavior, 61(1), 140–146. 10.1016/j.yhbeh.2011.11.008

Bhattacharyya, A., & Svendsen, C. (2003). Neurotrophins. In M. J. Aminoff & R. B. Daroff (Eds.), Encyclopedia of the Neurological Sciences (pp. 621–623). Academic Press. 10.1016/B0-12-226870-9/01013-3

Branchi, I., D’Andrea, I., Fiore, M., Di Fausto, V., Aloe, L., & Alleva, E. (2006). Early Social Enrichment Shapes Social Behavior and Nerve Growth Factor and Brain-Derived Neurotrophic Factor Levels in the Adult Mouse Brain. Biological Psychiatry, 60(7), 690– 696. 10.1016/j.biopsych.2006.01.005

Branchi, I., Francia, N., & Alleva, E. (2004). Epigenetic control of neurobehavioural plasticity: The role of neurotrophins. Behavioural Pharmacology, 15(5), 353–362. 10.1097/00008877-200409000-00006

Cardoso, S. D., Teles, M. C., & Oliveira, R. F. (2015). Neurogenomic mechanisms of social plasticity. Journal of Experimental Biology, 218(1), 140–149. 10.1242/jeb.106997

Chen, J., Kitanishi, T., Ikeda, T., Matsuki, N., & Yamada, M. K. (2007). Contextual learning induces an increase in the number of hippocampal CA1 neurons expressing high levels of BDNF. Neurobiology of Learning and Memory, 88(4), 409–415. 10.1016/j.nlm.2007.07.009

Chocyk, A., Bobula, B., Dudys, D., Przyborowska, A., Majcher-Maślanka, I., Hess, G., & Wędzony, K. (2013). Early-life stress affects the structural and functional plasticity of the medial prefrontal cortex in adolescent rats. European Journal of Neuroscience, 38(1), 2089– 2107. 10.1111/ejn.12208

Cirulli, F., Berry, A., & Alleva, E. (2003). Early disruption of the mother–infant relationship: Effects on brain plasticity and implications for psychopathology. Neuroscience & Biobehavioral Reviews, 27(1), 73–82. 10.1016/S0149-7634(03)00010-1

Clutton-Brock, T. H. (2006). Cooperative breeding in mammals. In Cooperation in Primates and Humans: Mechanisms and Evolution (pp. 173–190).

Coria-Lucero, C. D., Golini, R. S., Ponce, I. T., Deyurka, N., Anzulovich, A. C., Delgado, S. M., & Navigatore-Fonzo, L. S. (2016). Rhythmic Bdnf and TrkB expression patterns in the prefrontal cortex are lost in aged rats. BRAIN RESEARCH, 1653, 51–58. 10.1016/j.brainres.2016.10.019

D’Andrea, I., Alleva, E., & Branchi, I. (2007). Communal nesting, an early social enrichment, affects social competences but not learning and memory abilities at adulthood. Behavioural Brain Research, 183(1), 60–66. 10.1016/j.bbr.2007.05.029

Dey, C. J., Reddon, A. R., O’Connor, C. M., & Balshine, S. (2013). Network structure is related to social conflict in a cooperatively breeding fish. Animal Behaviour, 85(2), 395–402. 10.1016/j.anbehav.2012.11.012

Egan, M. F., Kojima, M., Callicott, J. H., Goldberg, T. E., Kolachana, B. S., Bertolino, A., Zaitsev, E., Gold, B., Goldman, D., Dean, M., Lu, B., & Weinberger, D. R. (2003). The BDNF val66met polymorphism affects activity-dependent secretion of BDNF and human memory and hippocampal function. Cell, 112(2), 257–269. 10.1016/S0092-8674(03)00035-7

Félix, A. S., & Oliveira, R. F. (2021). Integrative Neurobiology of Social Behavior in Cichlid Fish. In The Behavior, Ecology and Evolution of Cichlid Fishes (pp. 637–681). Springer Netherlands. 10.1007/978-94-024-2080-7_17

Fischer, S., Bessert-Nettelbeck, M., Kotrschal, A., & Taborsky, B. (2015). Rearing-Group Size Determines Social Competence and Brain Structure in a Cooperatively Breeding Cichlid. The American Naturalist, 186(1), 123–140. 10.1086/681636

Fischer, S., Bohn, L., Oberhummer, E., Nyman, C., & Taborsky, B. (2017). Divergence of developmental trajectories is triggered interactively by early social and ecological experience in a cooperative breeder. Proceedings of the National Academy of Sciences, 114(44), 201705934. 10.1073/pnas.1705934114

Gower, E. C. (1990). The long-term retention of events in monkey memory. Behavioural Brain Research, 38(3), 191–198. 10.1016/0166-4328(90)90174-D

Groenewoud, F., Frommen, J. G., Josi, D., Tanaka, H., Jungwirth, A., & Taborsky, M. (2016a). Predation risk drives social complexity in cooperative breeders. Proceedings of the National Academy of Sciences, 113(15), 4104–4109. 10.1073/pnas.1524178113

Groenewoud, F., Frommen, J. G., Josi, D., Tanaka, H., Jungwirth, A., & Taborsky, M. (2016b). Predation risk drives social complexity in cooperative breeders. Proceedings of the National Academy of Sciences, 113(15), 4104–4109. 10.1073/pnas.1524178113

Gustafsson, L., & Sutherland, W. J. (1988). The costs of reproduction in the collared flycatcher Ficedula albicollis. Nature, 335(6193), Article 6193. 10.1038/335813a0

Huang, E. J., & Reichardt, L. F. (2001). Neurotrophins: Roles in Neuronal Development and Function. Annual Review of Neuroscience, 24(1), 677–736. 10.1146/annurev.neuro.24.1.677

Isabel, G., Pascual, A., & Preat, T. (2004). Exclusive Consolidated Memory Phases in Drosophila. Science, 304(5673), 1024–1027. 10.1126/science.1094932

Jungwirth, A., Balzarini, V., Zöttl, M., Salzmann, A., Taborsky, M., & Frommen, J. G. (2019). Long-term individual marking of small freshwater fish: The utility of Visual Implant Elastomer tags. Behavioral Ecology and Sociobiology, 73(4), 1–11. 10.1007/S00265-019-2659-Y/FIGURES/3

Kaiser, S., Kruijver, F. P. M., Swaab, D. F., & Sachser, N. (2003). Early social stress in female guinea pigs induces a masculinization of adult behavior and corresponding changes in brain and neuroendocrine function. Behavioural Brain Research, 144(1), 199–210. 10.1016/S0166-4328(03)00077-9

Kamikubo, Y., Egashira, Y., Tanaka, T., Shinoda, Y., Tominaga-Yoshino, K., & Ogura, A. (2006). Long-lasting synaptic loss after repeated induction of LTD: Independence to the means of LTD induction. European Journal of Neuroscience, 24(6), 1606–1616. 10.1111/j.1460-9568.2006.05032.x

Kasumovic, M. M., & Brooks, R. C. (2011). It’s All Who You Know: The Evolution Of Socially Cued Anticipatory Plasticity As A Mating Strategy. The Quarterly Review of Biology, 86(3), 181–197. 10.1086/661119

Kempes, M. M., Gulickx, M. M. C., van Daalen, H. J. C., Louwerse, A. L., & Sterck, E. H. M. (2008). Social Competence Is Reduced in Socially Deprived Rhesus Monkeys (Macaca mulatta). Journal of Comparative Psychology, 122(1), 62–67. 10.1037/0735-7036.122.1.62

Kleim, J. A., & Jones, T. A. (2008). Principles of Experience-Dependent Neural Plasticity: Implications for Rehabilitation After Brain Damage. Journal of Speech, Language, and Hearing Research, 51(1), S225–S239. 10.1044/1092-4388(2008/018)

Klein, R., Nanduri, V., Jing, S. A., Lamballe, F., Tapley, P., Bryant, S., Cordon-Cardo, C., Jones, K. R., Reichardt, L. F., & Barbacid, M. (1991). The trkB tyrosine protein kinase is a receptor for brain-derived neurotrophic factor and neurotrophin-3. Cell, 66(2), 395–403. 10.1016/0092-8674(91)90628-c

Koenig, W. D., & Dickinson, J. L. (2016). Cooperative Breeding in Vertebrates: Studies of Ecology, Evolution, and Behavior. Cambridge University Press. https://books.google.ch/books?id=iCN0CwAAQBAJ

Kovalchuk, Y., Hanse, E., Kafitz, K. W., & Konnerth, A. (2002). Postsynaptic Induction of BDNF-Mediated Long-Term Potentiation. Science, 295(5560), 1729–1734. 10.1126/science.1067766

La Loggia, O., & Taborsky, B. (2024). Social competence is influenced by early but not late life social experience in a cooperatively breeding fish. Animal Behaviour, 213, 85–93. 10.1016/j.anbehav.2024.04.006

La Loggia, O., Wilson, A. J., & Taborsky, B. (2024). Early social complexity influences social behaviour but not social trajectories in a cooperatively breeding cichlid fish. Royal Society Open Science, 11(3), 230740. 10.1098/rsos.230740

Leal, G., Afonso, P. M., Salazar, I. L., & Duarte, C. B. (2015). Regulation of hippocampal synaptic plasticity by BDNF. Brain Research, 1621, 82–101. 10.1016/j.brainres.2014.10.019

Lee, H., Kim, D.-W., Remedios, R., Anthony, T. E., Chang, A., Madisen, L., Zeng, H., & Anderson, D. J. (2014). Scalable control of mounting and attack by Esr1+ neurons in the ventromedial hypothalamus. Nature, 509(7502), Article 7502. 10.1038/nature13169

Liedtke, J., & Schneider, J. M. (2017). Social makes smart: Rearing conditions affect learning and social behaviour in jumping spiders. Animal Cognition, 20(6), 1093–1106. 10.1007/s10071-017-1125-3

Liu, D., Diorio, J., Day, J. C., Francis, D. D., & Meaney, M. J. (2000). Maternal care, hippocampal synaptogenesis and cognitive development in rats. Nature Neuroscience, 3(8), Article 8. 10.1038/77702

Liu, Y., Donovan, M., Jia, X., & Wang, Z. (2019). The ventromedial hypothalamic circuitry and male alloparental behaviour in a socially monogamous rodent species. European Journal of Neuroscience, 50(11), 3689–3701. 10.1111/ejn.14550

Maejima, H., Kanemura, N., Kokubun, T., Murata, K., & Takayanagi, K. (2018). Exercise enhances cognitive function and neurotrophin expression in the hippocampus accompanied by changes in epigenetic programming in senescence-accelerated mice. Neuroscience Letters, 665, 67–73. 10.1016/J.NEULET.2017.11.023

McClellan, K. M., Parker, K. L., & Tobet, S. (2006). Development of the ventromedial nucleus of the hypothalamus. Frontiers in Neuroendocrinology, 27(2), 193–209. 10.1016/j.yfrne.2006.02.002

Minichiello, L., Korte, M., Wolfer, D., Kühn, R., Unsicker, K., Cestari, V., Rossi-Arnaud, C., Lipp, H.-P., Bonhoeffer, T., & Klein, R. (1999). Essential Role for TrkB Receptors in Hippocampus-Mediated Learning. Neuron, 24(2), 401–414. 10.1016/S0896-6273(00)80853-3

Mohr, R. A., Chang, Y., Bhandiwad, A. A., Forlano, P. M., & Sisneros, J. A. (2018). Brain Activation Patterns in Response to Conspecific and Heterospecific Social Acoustic Signals in Female Plainfin Midshipman Fish, Porichthys notatus. Brain, Behavior and Evolution, 91(1), 31–44. 10.1159/000487122

Murthy, S., & Gould, E. (2020). How Early Life Adversity Influences Defensive Circuitry. Trends in Neurosciences, 43(4), 200–212. 10.1016/j.tins.2020.02.001

Naguib, M., Nemitz, A., & Gil, D. (2006). Maternal developmental stress reduces reproductive success of female offspring in zebra finches. Proceedings of the Royal Society B: Biological Sciences, 273(1596), 1901–1905. 10.1098/rspb.2006.3526

Nyman, C., Fischer, S., Aubin-Horth, N., & Taborsky, B. (2017). Effect of the early social environment on behavioural and genomic responses to a social challenge in a cooperatively breeding vertebrate. Molecular Ecology, 26(12), 3186–3203. 10.1111/mec.14113

O’Connell, L. A., & Hofmann, H. A. (2011). The Vertebrate mesolimbic reward system and social behavior network: A comparative synthesis. The Journal of Comparative Neurology, 519(18), 3599–3639. 10.1002/cne.22735

O’Connell, L. A., & Hofmann, H. A. (2012). Evolution of a Vertebrate Social Decision-Making Network. Science, 336(6085), 1154–1157. 10.1126/science.1218889

Oliveira, R. F. (2009). Social behavior in context: Hormonal modulation of behavioral plasticity and social competence. Integrative and Comparative Biology, 49(4), 423–440. 10.1093/icb/icp055

Olivier, B. (1977). The ventromedial hypothalamus and aggressive behaviour in rats. Aggressive Behavior, 3(1), 47–56. 10.1002/1098-2337(1977)3:1<47::AID-AB2480030105>3.0.CO;2-H

Park, H., & Poo, M. M. (2013). Neurotrophin regulation of neural circuit development and function. Nature Reviews Neuroscience, 14(1), 7–23. 10.1038/nrn3379

Pfaffl, M. W. (2001). A new mathematical model for relative quantification in real-time RT– PCR. Nucleic Acids Research, 29(9), e45. 10.1093/nar/29.9.e45

Purves, D., Augustine, G. J., Fitzpatrick, D., Hall, W. C., LaMantia, A., McNamara, J. O., & Williams, S. M. (2004). Neuroscience. Third Edition. (D. Purves, G. J. Augustine, D. Fitzpatrick, W. C. Hall, A. LaMantia, J. O. McNamara, & S. M. Williams, Eds.). Sinauer Associates, Inc.

R Core Team. (2020). R: A language and environment for statistical computing. In R Foundation for Statistical Computing, Vienna, Austria. (p. URL https://www.R-project.org/) [Computer software]. http://www.mendeley.com/research/r-language-environment-statistical-computing-96/ papers2://publication/uuid/A1207DAB-22D3-4A04-82FB-D4DD5AD57C28

Razzoli, M., Domenici, E., Carboni, L., Rantamaki, T., Lindholm, J., Castrén, E., & Arban, R. (2011). A role for BDNF/TrkB signaling in behavioral and physiological consequences of social defeat stress. *Genes*, Brain and Behavior, 10(4), 424–433. 10.1111/J.1601-183X.2011.00681.X

Rodriguez-Tebar, A., Dechant, G., & Barde, Y.-A. (1990). Binding of brain-derived neurotrophic factor to the nerve growth factor receptor. Neuron, 4(4), 487–492. 10.1016/0896-6273(90)90107-Q

Sakuragi, S., Tominaga-Yoshino, K., & Ogura, A. (2013). Involvement of TrkB- and p75NTR-signaling pathways in two contrasting forms of long-lasting synaptic plasticity. Scientific Reports, 3(1), Article 1. 10.1038/srep03185

Segura, M., Pedreño, C., Obiols, J., Taurines, R., Pàmias, M., Grünblatt, E., & Gella, A. (2015). Neurotrophin blood-based gene expression and social cognition analysis in patients with autism spectrum disorder. Neurogenetics, 16(2), 123–131. 10.1007/S10048-014-0434-9

Shinoda, Y., Kamikubo, Y., Egashira, Y., Tominaga-Yoshino, K., & Ogura, A. (2005). Repetition of mGluR-dependent LTD causes slowly developing persistent reduction in synaptic strength accompanied by synapse elimination. Brain Research, 1042(1), 99–107. 10.1016/j.brainres.2005.02.028

Silva, B. A., Mattucci, C., Krzywkowski, P., Murana, E., Illarionova, A., Grinevich, V., Canteras, N. S., Ragozzino, D., & Gross, C. T. (2013). Independent hypothalamic circuits for social and predator fear. Nature Neuroscience, 16(12), 1731–1733. 10.1038/nn.3573

Soztutar, E., Colak, E., & Ulupinar, E. (2016). Gender- and anxiety level-dependent effects of perinatal stress exposure on medial prefrontal cortex. Experimental Neurology, 275, 274–284. 10.1016/j.expneurol.2015.06.005

Taborsky, B. (2017). Developmental Plasticity: Preparing for Life in a Complex World. In Advances in the Study of Behavior. 10.1016/bs.asb.2016.12.002

Taborsky, B. (2021). A positive feedback loop between sociality and social competence. Ethology, 127(10), 774–789. 10.1111/ETH.13201

Taborsky, B., Arnold, C., Junker, J., & Tschopp, A. (2012). The early social environment affects social competence in a cooperative breeder. Animal Behaviour, 83(4), 1067–1074. 10.1016/j.anbehav.2012.01.037

Taborsky, B., & Oliveira, R. F. (2012). Social competence: An evolutionary approach. Trends in Ecology and Evolution, 27(12), 679–688. 10.1016/j.tree.2012.09.003

Taborsky, B., Tschirren, L., Meunier, C., & Aubin-Horth, N. (2013). Stable reprogramming of brain transcription profiles by the early social environment in a cooperatively breeding fish. Proceedings of the Royal Society B: Biological Sciences, 280(1753), 20122605. 10.1098/rspb.2012.2605

Taborsky, M. (2016). Cichlid fishes: A model for the integrative study of social behavior. In W. D. Koenig & J. L. Dickinson (Eds.), Cooperative Breeding in Vertebrates (pp. 272–293). Cambridge University Press. 10.1017/CBO9781107338357.017

Teles, M. C., Cardoso, S. D., & Oliveira, R. F. (2016). Social plasticity relies on different neuroplasticity mechanisms across the brain social decision-making network in zebrafish. Frontiers in Behavioral Neuroscience, 10(FEB), 16. 10.3389/fnbeh.2016.00016

Thomason, M. E., Yoo, D. J., Glover, G. H., & Gotlib, I. H. (2009). BDNF genotype modulates resting functional connectivity in children. Frontiers in Human Neuroscience, 3, 55. 10.3389/neuro.09.055.2009

Tominaga-Yoshino, K., Kondo, S., Tamotsu, S., & Ogura, A. (2002). Repetitive activation of protein kinase A induces slow and persistent potentiation associated with synaptogenesis in cultured hippocampus. Neuroscience Research, 44(4), 357–367. 10.1016/S0168-0102(02)00155-4

Tominaga-Yoshino, K., Urakubo, T., Okada, M., Matsuda, H., & Ogura, A. (2008). Repetitive induction of late-phase LTP produces long-lasting synaptic enhancement accompanied by synaptogenesis in cultured hippocampal slices. Hippocampus, 18(3), 281–293. 10.1002/hipo.20391

Trutti, A. C., Mulder, M. J., Hommel, B., & Forstmann, B. U. (2019). Functional neuroanatomical review of the ventral tegmental area. NeuroImage, 191, 258–268. 10.1016/j.neuroimage.2019.01.062

Watve, M., & Taborsky, B. (2019). Presence of parents during early rearing affects offspring responses towards predators. Animal Behaviour, 158, 239–247. 10.1016/j.anbehav.2019.09.012

Zanin, J. P., Montroull, L., Volosin, M., & Friedman, W. (2019). The p75 Neurotrophin Receptor Facilitates TrkB Signaling and Function in Rat Hippocampal Neurons. Frontiers in Cellular Neuroscience. 10.3389/fncel.2019.00485

